# Pupil correlates of decision variables in mice playing a competitive mixed-strategy game

**DOI:** 10.1101/2021.08.05.455292

**Authors:** Hongli Wang, Heather K. Ortega, Huriye Atilgan, Cayla E. Murphy, Alex C. Kwan

**Affiliations:** Interdepartmental Neuroscience Program, Yale University School of Medicine, New Haven, Connecticut, 06511, USA; Department of Psychiatry, Yale University School of Medicine, New Haven, Connecticut, 06511, USA; Department of Neuroscience, Yale University School of Medicine, New Haven, Connecticut, 06511, USA

## Abstract

In a competitive game involving an animal and an opponent, the outcome is contingent on the choices of both players. To succeed, the animal must continually adapt to competitive pressure, or else risk being exploited and lose out on rewards. In this study, we demonstrate that head-fixed mice can be trained to play the iterative competitive game ‘matching pennies’ against a virtual computer opponent. We find that the animals’ performance is well described by a hybrid computational model that includes Q-learning and choice kernels. Comparing between matching pennies and a non-competitive two-armed bandit task, we show that the tasks encourage animals to operate at different regimes of reinforcement learning. To understand the involvement of neuromodulatory mechanisms, we measure fluctuations in pupil size and use multiple linear regression to relate the trial-by-trial transient pupil responses to decision-related variables. The analysis reveals that pupil responses are modulated by observable variables, including choice and outcome, as well as latent variables for value updating, but not action selection. Collectively, these results establish a paradigm for studying competitive decision-making in head-fixed mice and provide insights into the role of arousal-linked neuromodulation in the decision process.

## INTRODUCTION

Animals learn from the outcomes of their past actions. The decision-making process can be casted in the framework of reinforcement learning (Sutton and Barto, 1998), which provides a quantitative approach to characterize how animals choose among multiple options based on prior experience. This approach, when applied to rodents and combined with powerful molecular, genetic, electrophysiological, and imaging methods, has yielded novel insights into the neural circuits involved in reward-based learning (Bari et al., 2019; Groman et al., 2019; Hattori et al., 2019; Ito and Doya, 2009; Sul et al., 2011; Sul et al., 2010; Tai et al., 2012). To date most studies in rodents relied on tasks involving a pure strategy, where there is always a particular action that is optimal in a given situation. However, pure strategies are not always available. For instance, in a two-player game (e.g., rock-paper-scissors) where the outcome depends on both the animal’s and an opponent’s choices, pure strategies are inadequate because an opponent can predict tendencies and exploit to win. Instead, the animal should adopt a mixed strategy, in which two or more pure strategies are chosen probabilistically. Analyses of behavior under competitive pressure fall into the purview of game theory (Camerer, 2003), and provide a unique window into the social and adaptive aspects of decision-making (Lee, 2008).

Research in humans and non-human primates has identified numerous brain regions contributing to decision-making during two-player games. For example, a functional imaging study in humans showed widespread representation of reward signals in the brain during multiple types of games against computerized opponents (Vickery et al., 2011). Signals related to choices and the mentalization of an opponent’s actions were localized to several brain regions including the prefrontal cortex (Hampton et al., 2008; Vickery et al., 2011). Electrophysiological recordings in macaque monkeys demonstrated spiking activity patterns that suggest neural representations of various decision variables in prefrontal cortical regions, lateral intraparietal cortex, and amygdala (Barraclough et al., 2004; Chang et al., 2013; Dal Monte et al., 2020; Haroush and Williams, 2015; Seo et al., 2007, 2009). However, there have only been a few reports of rodents engaging in two-player games (Tervo et al., 2014; Wood et al., 2016), and the associated neural correlates are less clear.

The neuromodulator norepinephrine (NE) may have an important role for performance during two-player games. Prior literature has linked central noradrenergic tone to behavioral flexibility (Aston-Jones and Cohen, 2005; Bouret and Sara, 2005). In one pioneering study, Tervo et al. taught rats to play a matching pennies game with a computer opponent (Tervo et al., 2014). By manipulating neural activity in locus coeruleus (LC), they showed that elevating central NE tone can suppress firing in the anterior cingulate cortex, which in turn reduces the influence of reinforcement history and promotes stochastic behavior. A different way to study neuromodulatory tone is to measure pupil size, which is often treated as a readout of NE levels in the neocortex (Gilzenrat et al., 2010; Joshi et al., 2016; Reimer et al., 2016). Studies of pupillary dynamics during tasks further support the idea that NE is important for flexible decisions. For instance, the baseline pupil size is shown to correlate with biases in the explore-exploit trade-off and attentional set shifting (Jepma and Nieuwenhuis, 2011; Pajkossy et al., 2017). The trial-by-trial transient change in pupil size is reported to associate with many task-relevant variables including upcoming choice, expected outcome, values of the choices, and uncertainties (de Gee et al., 2014; Hess and Polt, 1960; Qiyuan et al., 1985; Van Slooten et al., 2018). Given these prior results, it seems likely that pupil fluctuations can be leveraged to study the neuromodulatory mechanisms underlying adaptive action selection during competitive two-player games.

We have two goals for the current study. First, we want to know if head-fixed mice can compete proficiently in a two-player game. Second, we want to characterize pupil fluctuations to gain insights into the role of neuromodulation in decision-making under competitive pressure. To this end, we designed a behavioral paradigm for a head-fixed mouse to play iterative matching pennies against a computer-controlled virtual opponent. We found that mice can perform at a level close to the optimal reward rate by exhibiting choice behavior consistent with a mix of reinforcement learning and choice perseveration. We underscored the unique choice behavior during matching pennies, by comparing with performance in the more widely used two-armed bandit task. Finally, we measured within-trial changes in pupil size, and showed that the transient pupillary responses are associated with the choice, outcome, and latent variables for value updating.

## MATERIALS AND METHODS

### Animals

All animal procedures were conducted in accordance with procedures approved by the Institutional Animal Care and Use Committee at Yale University. Adult male C57BL/6J mice (>P56; #000664, Jackson Laboratory) were used for all experiments. Mice were housed in groups of 3-5 animals with 12-hr-dark/12-hr-light cycle control (lights off at 19:00).

### Surgical Procedures

Anesthesia was induced with 2% isoflurane in oxygen before the surgery. The isoflurane was lowered to 1–1.2% during the surgical procedures. The mouse was placed on a water-circulating heating pad (TP-700, Gaymar Stryker) in a stereotaxic frame (David Kopf Instruments). After injecting carprofen (5 mg/kg, s.c.; #024751, Butler Animal Health) and dexamethasone (3 mg/kg, s.c.; Dexaject SP, #002459, Henry Shein Animal Health), the scalp was removed to expose the skull. A custom-made stainless-steel head plate (eMachineShop) was glued onto the skull with MetaBond (C&B, Parkell, Inc.). Carprofen (5 mg/kg, s.c.) was injected each day for the following three days. Mice were given 7 days to recover from the surgery before any behavioral training.

### Behavioral setup

The training apparatus was based on a previous design from our prior studies (Siniscalchi et al., 2016; Siniscalchi et al., 2019). Detailed instruction to construct the apparatus is available at https://github.com/Kwan-Lab/behavioral-rigs. The mouse with a head plate implant was head-fixed to a stainless-steel holder (eMachineShop). The animal sat inside an acrylic tube (8486K433; McMaster-Carr), which limited gross movements though allowed postural adjustments. A lick port with two lick spouts was positioned in front of the subject. The spouts were constructed with blunted 20-gauge stainless-steel needles. Contact with the lick spout – which was how the animal indicated its choices – was detected through wires that were soldered onto the spout and a battery-powered lick detection electronic circuit. Output signals from the circuit were sent to a computer via a data acquisition unit (USB-201, Measurement Computing) and logged by the Presentation software (Neurobehavioral Systems, Inc.). Water delivery from the lick spouts was controlled independently for each spout by two solenoid fluid valves (MB202-V-A-3-0-L-204; Gems Sensors & Controls). The amount of water was calibrated to ~4 μL per pulse by adjusting the duration of the electrical pulse sent by the Presentation software via a second data acquisition unit (USB-201, Measurement Computing). Two speakers (S120, Logitech) were placed in front of the mouse to play the sound cue. The whole setup was placed inside an audio-visual cart with walls lined with soundproof acoustic foams (5692T49, McMaster-Carr). A monochrome camera (GigE G3-GM11-M1920, Dalsa) with a 55 mm telecentric lens (TEC-55, Computar) was aimed at the right eye. Video was acquired at 20 Hz. A dimmable, white light source (LT-T6, Aukey) was used to provide ambient light, such that the baseline pupil size was moderate and fluctuations around the baseline was detectable. The computer running the Presentation software would send TTL pulses to a computer controlling the camera through a USB data acquisition device (USB-201; Measurement Computing). The camera-connected computer would run a custom script written in MATLAB 9.7 (MathWorks) that logged the timing of the TTL pulses so that the behavioral log files generated by the Presentation software could be aligned to the video recordings. In a small subset of experiments, we captured videos from both left and right eyes by mounting two identical camera systems on both sides of the animal.

### Behavioral training – Matching pennies

All of the procedures for initial shaping as well as the final matching pennies task were written using the scripting language in the Presentation software. The animals were fluid-restricted. Water was provided during the one behavioral session daily. On the days when the animals were not trained (typically 1 day a week), a water bottle was placed in the home cage for 5 min of ad libitum consumption. All animals underwent two shaping phases before training. For phase 1 (2 days), the animals were habituated to the apparatus. They may lick either spout. A water reward would be delivered for every lick in the corresponding spout, as long as a minimum of 1 s has occurred since the last reward. The session would terminate after the animals acquired 100 rewards. For phase 2 (~4 weeks), the animals were introduced to the trial structure and learned to suppress impulsive licks. At the start of each trial, a 5 kHz sound cue lasting for 0.2 s was played. From the onset of the sound cue, the mouse had a window of 2 s to make a response by licking either of the spouts. If a lick was detected during the response window, a water reward would be delivered in the corresponding spout and there was a fixed 3 s period for consumption following the lick. If no lick was detected, the fixed 3 s consumption window would still be presented following the end of the response window. From the end of the consumption window, an inter-trial interval (ITI) began. The duration of the ITI in seconds was drawn from a truncated exponential distribution with λ = 1/3 and boundaries of 1 and 5. If the animal emitted one or more licks during the ITI, then additional time drawn again from the same distribution would be appended to the ITI. If the mouse licked again during the appended time, yet another additional time would be appended, up to a total of 5 draws including the initial draw. When the ITI ended, a new trial would begin. This trial timing was the same as what would be used for matching pennies. Particularly, the goal for the shaping was to habituate and introduce lick suppression. Although the mouse could theoretically get water from either spout, the animal tended to favor heavily one spout during the shaping procedures. The animal would advance to playing the matching pennies game when the average number of ITI draws per trial was lower than 1.2 for three consecutive sessions.

For matching pennies, the mouse played against a virtual opponent in the form of a computer agent. At the start of each trial, the agent made a choice (left or right). If the mouse selected the same choice as the computer, a water reward would be delivered in the corresponding spout. Otherwise, no reward was presented. Importantly, the computer agent was designed to provide competitive pressure by acting according to prediction of the animal’s choices. Specifically, it was programmed to be the same as ‘algorithm 2’ (Barraclough et al., 2004; Lee et al., 2004) or ‘competitor 1’ (Tervo et al., 2014) in previous studies. Briefly, the agent had a record of the mouse’s entire choice and reward history within the current session. The agent calculated the conditional probabilities that the animal would choose left given sequences of the preceding N choices (N = 0 – 4) and sequences of preceding N choice-outcome combinations (N = 1 – 4). The binomial test was used to test each of the 9 conditional probabilities against the null hypothesis that the mouse would choose left with a probability of 0.5. If none of the null hypothesis was rejected, the agent would randomly choose either target with equal probabilities. If one or more hypotheses were rejected, the agent would generate the counter choice with the statistically significant conditional probability that was farther away from 0.5. A session would terminate automatically when no response was logged for 10 consecutive trials. When an animal reached a 40% reward rate for 3 consecutive sessions (~4 weeks), then its performance was considered stable and the subsequent sessions were included in the following analysis.

### Behavioral training – Two-armed bandit

We used the same training apparatus and the same scripting language in the Presentation software to program the two-armed bandit task, which had the same trial timing as matching pennies. The shaping procedures relied on similar tactics, but the details were different, involving three shaping phases before training. For phase 0 (1 day), the experimenter manually delivered 50 water rewards through each port (for a total of 100 rewards) and monitored for consistent licking. A 5 kHz sound cue lasting for 0.2 s was played at the same time as water delivery. If the animal was not licking to consume the water rewards, the experimenter used a blunted syringe to guide the animal to lick the spout. For phase 1 (1 day), the animal was introduced to the trial structure and learned to suppress impulsive licks. At the start of each trial, A 5 kHz sound cue lasting for 0.2 s was played. From the onset of the sound cue, the mouse had a window of 5 s to make a response by licking either of the spouts. If a lick was detected during the response window, a water reward would be delivered from the corresponding spout and there was a fixed 3 s period for consumption following the lick. If no lick was detected, no reward was given during the consumption period. Next, an ITI began. The duration of the ITI in seconds was drawn from a truncated exponential distribution with λ = 1/3 and boundaries of 1 and 5. If the animal emitted one or more licks during the ITI, then additional time drawn again from the same truncated exponential distribution would be appended to the ITI. If the mouse licked again during the appended time, yet another additional time would be appended, up to a total of 5 draws including the initial draw. When the ITI ended, a new trial would begin. The session ends after the animal accumulates 100 rewards. In particular, the goal for phase 1 was to habituate and introduce lick suppression. For phase 2, the animal continued to lick following the cue for a water reward in a similar fashion to phase 1. However, the response window was shortened to 2 s, meaning the animal had only 2 s to lick following the cue to receive reward. The trial timing with shortened response window was the same as what would be used for the two-armed bandit task. Moreover, the animal must alternate sides (choose left if the last reward came from a right lick, and vice versa). They would not be rewarded for choosing the same spout repeatedly to discourage the development of a side bias. The animal could proceed to the final task if they achieved 200 rewards in a single session.

For the two-armed bandit task, the two options (left and right) were associated with different reward probabilities. In each trial, when an animal chose an option, a reward was delivered stochastically based on the assigned reward probability. In our implementation, there were two sets of reward probabilities: 0.7:0.1 and 0.1:0.7. A set of reward probabilities was maintained across a block of trials. Within a block, once the mouse had chosen the option with high reward probability (hit trials) for ten times, then on any given trial there was a probability of 1/11 that the reward probabilities would change. Thus, the number of trials in a block after ten hits followed a geometric distribution with *μ* = 11. There were no explicit cues to indicate a block switch, therefore the animal had to infer the current situation through experience. A session would terminate automatically when no response was logged for 20 consecutive trials. Animals are considered proficient when they chose the spout with higher reward probability on at least 50% of trials on 3 consecutive sessions.

### Preprocessing of behavioral data

A total of 115 sessions from 13 mice were included in the study. For matching pennies, data came from 81 sessions from 5 mice. For two-armed bandit, data came from 34 sessions from 8 mice, including 26 sessions from 4 mice with single-pupil recording and 8 sessions from the other 4 mice with double-pupil recording. All of the sessions contained both behavioral and pupillometry data. For behavior, the log file saved by the Presentation software contained timestamps for all events that occurred during a session. Analyses of the behavioral data were done in MATLAB. For matching pennies, towards the end of each session, the animals tended to select the same option for around 30 trials before ceasing to respond. To avoid these repetitive trials in the analyses, for each session, the running 3-choice entropy (see below) of a 30-trial window was calculated, and the MATLAB function *ischange* was used to fit with a piecewise linear function. The trial when the fitted function fell below a value of 1 was identified as the ‘last trial’, and all subsequent trials were discarded for the analysis. In cases if the performance recovered after the detected last trial to a value greater than 1, or if the fitted function did not fall below a value of 1 and no ‘last trial’ was detected, the entire session was used for analysis.

### Analysis of behavioral data – Entropy

To quantify the randomness in the animals’ choices, the 3-choice entropy of the choice sequence is calculated by:

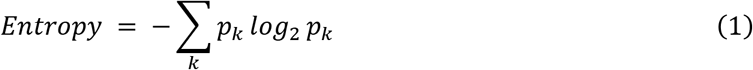

where *p_k_* is the frequency of occurrence of a 3-choice pattern in a session. Because there were 2 options to choose from, there were 2^3^ = 8 potential patterns possible. The maximum value for entropy is 3 bits.

### Analysis of behavioral data – Computational models

To quantify the choice behavior, we considered 5 computational models. The primary model used in the paper is a Q-learning with forgetting model plus choice kernels (FQ_RPE_CK) (Katahira, 2015; Wilson and Collins, 2019). On trial *n*, for a choice *c_n_* that leads to an outcome *r_n_*, the action value 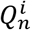, associated with an action *i* is updated by:

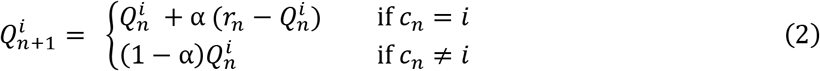

where α is the learning rate, and the forgetting rate for the unchosen action. In our task, there are two options, so *i* ∈ {*L, R*}. For the outcome, *r_n_* = 1 for reward, 0 for no reward. Moreover, to capture the animal’s tendency to make decisions based purely on previous choices, there are choice kernels 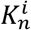 updated by:

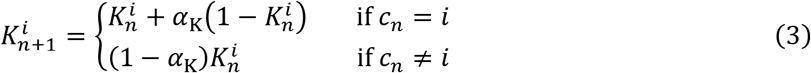

where *α*_K_ is the learning rate of the choice kernel. For action selection, the probability to choose action *i* on trial *n* is given by a softmax function:

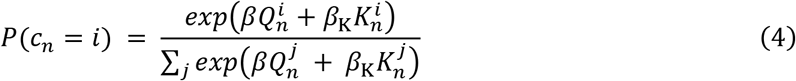

where *β* and *β*_K_ are the inverse temperature parameters for action values and choice kernels respectively.

We compared the FQ_RPE_CK model against 4 other models. For the win-stay-lose-switch model (WSLS), the probability to choose action *i* on trial *n* is given by:

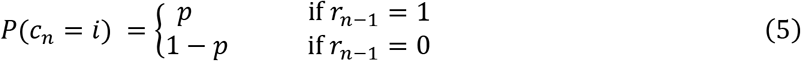

where *p* is the probability that the animal followed the win-stay-lose-switch strategy.

For the Q-learning model (Q_RPE), the action value is updated by:

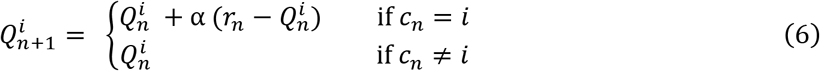

and the probability to choose action *i* at trial *n* is then given by:

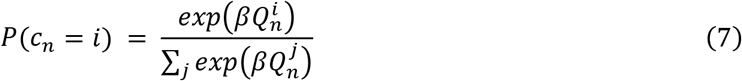

For the forgetting Q-learning model (FQ_RPE), the action values are updated by equation (2), and the probability to choose action *i* on trial *n* is given by equation (7).

For the differential Q-learning model (DQ_RPE) (Caze and van der Meer, 2013; Katahira, 2018), the action value is updated by:

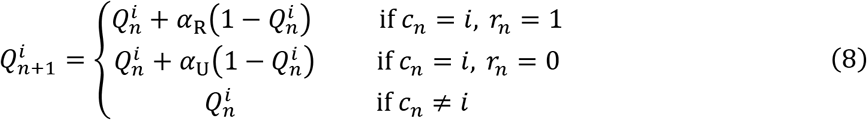

where *α*_R_ and *α*_U_ are the learning rates for rewarded and unrewarded trials, respectively. The probability to choose action *i* on trial *n* is given by equation (7).

### Analysis of behavioral data – Model fitting and comparison

To fit the computational models to the behavioral data, for each subject, the sequence of choices and outcomes were concatenated across sessions. Each model was fitted to the data using a maximum-likelihood algorithm implemented with the *fmincon* function in MATLAB, with the constraints 0 ≤ *α, α*_K_, *α*_R_, *α*_U_ ≤ 1, 0 < *β,β*_K_, and 0 ≤ *p* ≤ 1. These fits also yielded latent decision variables such as action values and choice kernels that would be used for the subsequent multiple linear regression analyses. For model comparison, we calculated the Bayesian information criterion (BIC) for each of the model fits.

### Preprocessing of pupillometry data

To extract the coordinates of the pupil from the video recordings, we used DeepLabCut (DLC) 2.0 (Mathis et al., 2018; Nath et al., 2019), ran on Jupyter Notebook on Google’s cloud servers. A small subset of the video frames was manually analyzed, with the experimenter annotating 5 labels including the central, uppermost, leftmost, lowermost, and rightmost points of the pupil. The annotated frames were fed to DLC to train a deep neural network, which analyzed the remainder of the video to produce the 5 labels. From the labels, the pupil diameter was computed by taking the distance between the leftmost and rightmost labels. We did not use the other labels, because the estimates of the lowermost points were unstable, sometimes jumping in consecutive frames due to interference from the lower eyelid. The pupil diameter signal was further processed through a 4 Hz lowpass filter with the MATLAB function *lowpass*. Then any frames with outliers that were greater than 3 scaled median absolute deviation (MAD) from the median were deleted using the MATLAB function *isoutlier*. Using a 10-minute moving window to account for drift over a session, the signal was converted to z-score. Finally, we calculated the pupil response for each trial by subtracting the instantaneous z-score from −1 to 5 s relative to cue onset by the baseline z-score, which was defined as the mean z-score from −2 to −1 s relative to cue onset.

### Analysis of pupil data – Multiple linear regression

To determine how pupil responses may relate to choices and outcomes, we used multiple linear regression:

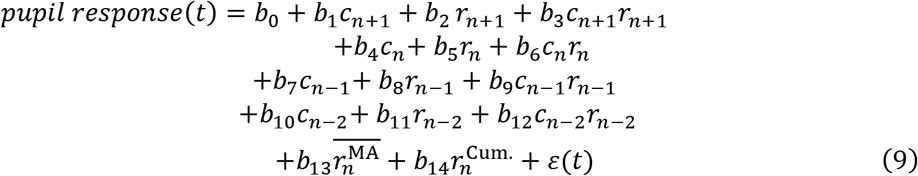

where *pupil response*(*t*) is the pupil response at time *t* in trial *n*, *c*_*n*+1_, *c_n_*, *c*_*n*-1_, *c*_*n*-2_ are the choices made on the next trial, the current trial, the previous trial, and the trial before the previous trial, respectively, *r*_*n*+1_, *r_n_*, *r*_*n*-1_, *r*_*n*-2_ are the outcomes for the next trial, the current trial, the previous trial, and the trial before the previous trial, respectively, *b*_0_,…, *b*_14_ are the regression coefficients, and *ε*(*t*) is the error term. Choices were dummy-coded as 0 for left responses and 1 for right responses. Outcomes were dummy-coded as 0 for no-reward and 1 for reward. For the last 2 predictors in Eq. 9, 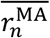 is the average reward over the previous 20 trials, given by the equation:

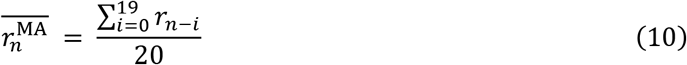

The term 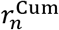 indicates the normalized cumulative reward during the session, calculated by:

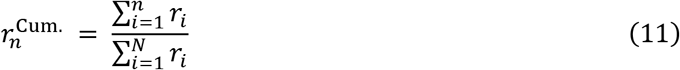

where *n* denotes the current trial number and *N* is the total number of trials in the session.

To determine how pupil responses may relate to latent decision variables for action selection, we used multiple linear regression:

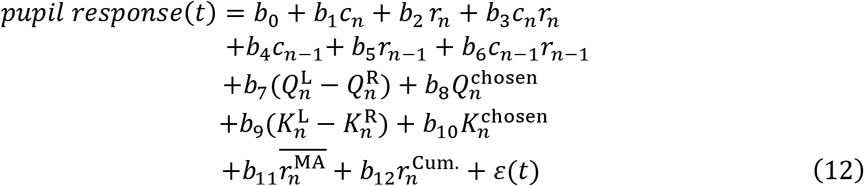

where 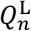 and 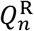 denote the action values of the left and right choices in trial *n*, respectively, 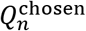 is the value of the action chosen in trial *n*, 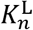 and 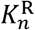 are the choice kernels of the left and right choices in trial *n*, respectively, 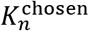 is the choice kernel of the action chosen in trial *n*.

To determine how pupil responses may relate to latent decision variables for value updating, we used multiple linear regression, adapting the equation from (Sul et al., 2010):

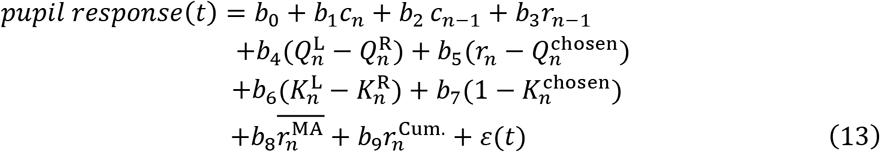

where 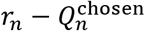 is the reward prediction error (RPE), and 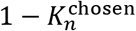 is the error term used to update the choice kernels, or choice kernel error (CKE).

For each session, the regression coefficients were determined by fitting the equations to data using the MATLAB function *fitlm*. The fit was done in 100-ms time bins that span from −3 to 5 s relative to cue onset, using mean pupil response within the time bins. To summarize the results, for each predictor, we calculated the proportion of sessions in which the regression coefficient was different from zero (*P* < 0.01). To determine if the proportion was significantly different from chance, we performed a chi-square test against the null hypothesis that there was a 1% probability that a given predictor was mischaracterized as significant by chance in a single session.

### Code accessibility

The data and code that support the findings of this study will be made publicly available at http://github.com/Kwan-Lab.

## RESULTS

### Mice played matching pennies against a computer opponent

We trained head-fixed mice to play matching pennies against a computer opponent (***Figure 1A***). In this iterative version of matching pennies, each trial the mouse and the computer would choose left or right. The mouse received a water reward only if the actions matched. The corresponding payoff matrix is shown in ***Figure 1B***. This game was challenging for the mouse because the computer opponent had access to the complete history of choices and rewards over the session, and was programmed to predict the mouse’s next choice in order to make the counter action (see Materials and Methods; same as ‘algorithm 2’ in (Lee et al., 2004) and ‘competitor 1’ in (Tervo et al., 2014)). ***Figure 1C*** shows the trial structure. A 0.2 s, 5 kHz sound cue indicated the start of the trial. Within a 2-s response window, the mouse could make a directional tongue lick to the left or right spout to indicate its choice. Based on the animal’s and computer’s choices and the payoff matrix, the animal might receive a water reward after the response. To minimize pre-cue licks, the inter-trial interval was drawn from a truncated exponential distribution and would extend if the mouse could not suppress licking (see Materials and Methods). Mice were trained daily and, on average, took about 4 weeks to follow the trial timing to suppress pre-cue licking, and then another 4 weeks playing matching pennies to achieve a performance threshold of 40% reward rate for 3 consecutive sessions. All the data and analyses presented in the paper came from sessions after the threshold was attained.

**Figure 1.**
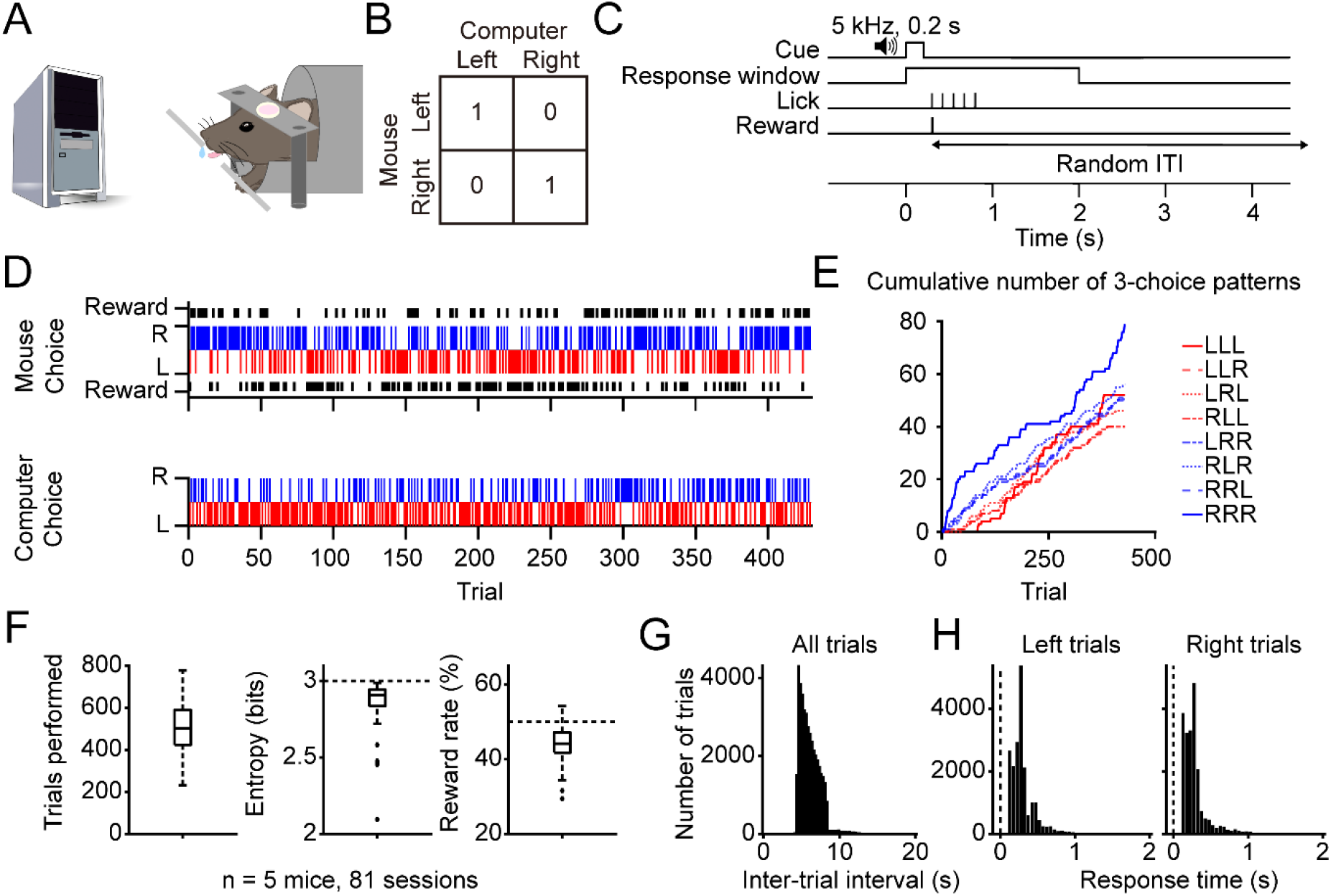
Performance of head-fixed mice in a matching pennies game. **(A)** A schematic illustration of the competitive game. Head-fixed mouse makes a left or right choice by licking the spouts. A computer monitors the mouse’s past choices and outcomes, generating its own left or right choice every trial. **(B)** The payoff matrix of the game. The mouse receives a water reward if it chooses the same choice as the computer does in the same trial. **(C)** Trial timing: the mouse waits for a go cue, licks a spout to indicate its response, and the outcome is delivered immediately. A random inter-trial interval follows the outcome. **(D)** An example session. The reward rate for this session was 52.1%. Top: the mouse’s choices and outcomes. Bottom: the computer’s choices. Blue and red bars indicate right and left choices respectively. Black bars indicate rewards. **(E)** Cumulative number of different 3-choice patterns detected as the mouse progressed in the session shown in (D). **(F)** Summary from 81 sessions. Left: the average trials performed each session is 513±13. Middle: the average entropy of the 3-choice sequences is 2.87±0.02. Right: the average reward rate is 44.0%±0.5%. **(G)** The histogram of the inter-trial interval durations for all trials. Inter-trial interval would range from 4 to 8 s-long if the mouse did not lick to trigger additions to the inter-trial interval. **(H)** The response times for trials in which the mouse chose left (left) or right (right).

The dataset for matching pennies included 81 behavioral sessions with concurrent pupil measurements (n = 5 mice). For matching pennies, the Nash equilibrium indicates that rational players should choose left and right with equal probabilities, which would yield a long-run reward rate of 50%. In an example session, plotting the choices and rewards for a mouse and the choices for the computer showed that the animal indeed exhibited a great degree of stochasticity in its choice pattern (***Figure 1D***). This could be seen more clearly by looking at the cumulative occurrences of various plausible 3-choice sequences (***Figure 1E***). Although there was sometimes a slight preference for certain patterns, such as for right-right-right early in this example session, throughout this session the animal continually employed different choice sequences. On average, animals performed 513±13 trials per session (mean±s.e.m.; ***Figure 1F***). The entropy for 3-choice sequences, a measure of the stochasticity in choices, was 2.87±0.02 bits, close to the theoretical upper bound of 3 bits. Consequently, the computer was only mildly effective at predicting the mouse’s choices, and mice earned an average reward rate of 44.0±0.5%. We note that although the reward rate compared favorably with the optimal reward rate of 50%, the difference was statistically significant (*P* = 1.4 x 10^-20^, one-sample one-tailed t-test). For the inter-trial interval, 98.4% of the trials fell within the truncated exponential distribution used to produce the random durations, and only 1.6% of the trials forming a small tail reflecting the extended duration triggered by a pre-cue impulsive lick (***Figure 1G***). Response times were stereotypical and similar for the left and right spouts (***Figure 1H***). These inter-trial interval and response time analyses demonstrate effective pre-cue lick suppression and that the animal makes the first physical indication of its choice after cue onset. Altogether, the results show that head-fixed mice can play matching pennies against a computer opponent. Given that humans and macaques likewise play matching pennies imperfectly and not at Nash equilibrium (Erev and Roth, 1998; Lee et al., 2004), here we found that reward rate was decent but also suboptimal for mice, suggesting that the animals had certain residual tendencies that were exploited by the computer opponent.

### Animals’ behavior was captured by a hybrid model with reinforcement learning and choice kernels

What is the strategy that characterizes the tendencies in the animal’s behavior? Previous studies in macaque monkeys found that reinforcement learning can account in part for the animals’ behavior in matching pennies (Lee et al., 2004; Seo et al., 2007). We therefore compared between a range of strategies: win-stay-lose-switch (WSLS), and reinforcement learning algorithms including Q-learning (Q_RPE), differential Q-learning (DQ_RPE), and forgetting Q-learning (FQ_RPE) (Ito and Doya, 2009) (see Materials and Methods). Fitting each model to the behavioral data, the Bayesian information criterion (BIC) values indicated that FQ_RPE was most consistent with the choice behavior of the mice. To further improve the model, we noted that mice exhibited serial choice dependence and sometimes favored picking the same choice in successive trials (e.g., ***Figure 1E***). To capture perseverative behavior, we added choice kernels (Wilson and Collins, 2019) to create the FQ_RPE_CK algorithm. ***Figure 2A*** illustrates graphically the FQ_RPE_CK scheme (henceforth called the ‘hybrid model’), with reinforcement learning through FQ_RPE and perseveration through choice kernels. Note that both the action values and choice kernels were updated on a trial-by-trial basis, and each had its own learning rate (*α* and *α*_K_) and inverse temperature parameters (*β* and *β*_K_; see Materials and Methods). A comparison of all 5 models indicated that the hybrid model provided the most accurate fit to the behavioral data (***Figure 2B***). ***Figure 2C*** shows a representative session in which we plotted the animal’s choices and outcomes along with the latent variables, including action values and choice kernels, estimated by the hybrid model. The model-estimated probability of choosing left (black line, ***Figure 2C***) tracked the actual choice pattern (gray line, ***Figure 2C***). Taken together, these analyses indicate that the mouse’s behavior during matching pennies can be quantified using a hybrid model including reinforcement learning and choice kernels. For the remainder of the analyses, we will quantify the animal’s strategy using the hybrid model.

**Figure 2.**
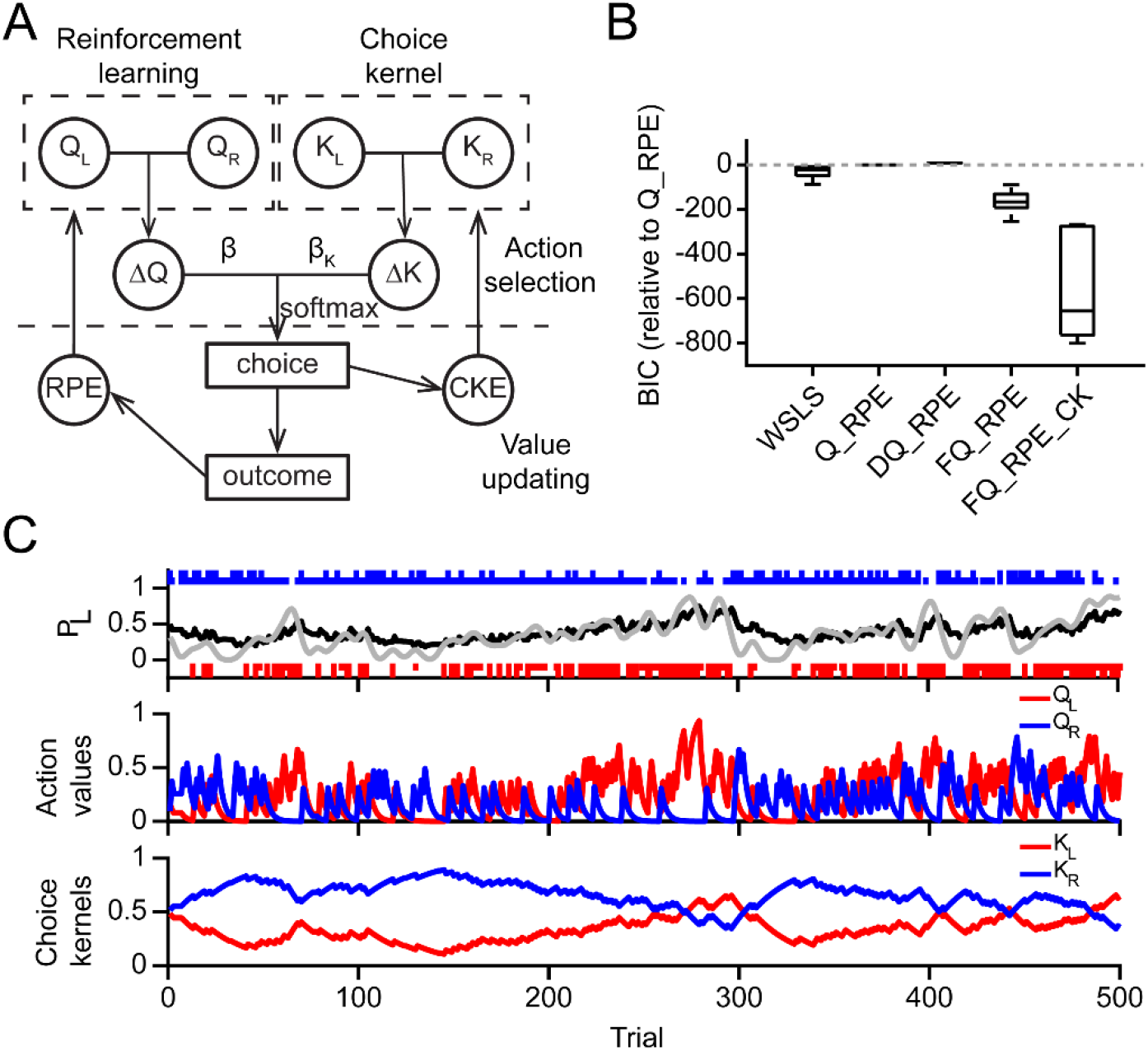
Computational modeling of the animals’ behavior during matching pennies. **(A)** Schematic of the hybrid FQ_RPE_CK model. *Q*_L_, *Q*_R_ denote the action values for left and right choices; *K*_L_, *K*_R_ denote the choice kernels for left and right choices. The agent chooses based on a weighted sum of action-value and choice-kernel differences (Δ*Q*, Δ*K*). The chosen action is used to compute the choice kernel error (CKE), which is used to update the choice kernels. The outcome is used to calculate the reward prediction error (RPE), which is used to update the action values. **(B)** Model comparison using BIC. **(C)** An example of the time course of the latent variables and predicted behavior. Top: Long red and blue bars indicate rewarded left and right choices; short red and blue bars indicate unrewarded left and right choices. Gray line shows the observed probability to choose left, smoothed by a gaussian kernel. Black line shows the probability to choose left predicted by the hybrid model. Middle: the action values of left (red) and right (blue) choices estimated by the hybrid model. Bottom: the choice kernels of left (red) and right (blue) choices estimated by the hybrid model.

### Learning strategy in matching pennies in contrast to two-armed bandit task

To gain further insights into how the mouse competes against a computer opponent, we compared performance between matching pennies and the two-armed bandit task, which is a popular, non-competitive paradigm for assaying reward-based learning in rodents. In our implementation of the two-armed bandit, trials were organized into blocks, and each block was associated with one of two sets of reward probabilities (top panel, ***Figure 3A***). The block switched randomly (see Materials and Methods) so that the mouse could not predict the switch with certainty and must infer based on past choices and outcomes. Moreover, the timing of each trial was design to be exactly the same between matching pennies and two-armed bandit (***Figure 1C***). The bottom panel of ***Figure 3A*** presents an example session of a mouse engaging in two-armed bandit, showing that the mouse readily adapting its actions in response to the reversing reward probabilities. The same hybrid FQ_RPE_CK model could be used to quantify the behavior, with the estimated choice probability tracking the mouse’s choice pattern (***Figure supplement 1***). Overall, mice performed the task well, averaging 702±26 trials per session with a 43.5±1.1% reward rate (n = 26 sessions from 4 mice; ***Figure 3B***). Model comparison based on BIC indicated that the hybrid model outperformed win-stay lose-switch and other Q-learning-based algorithms (***Figure 3C***).

**Figure 3.**
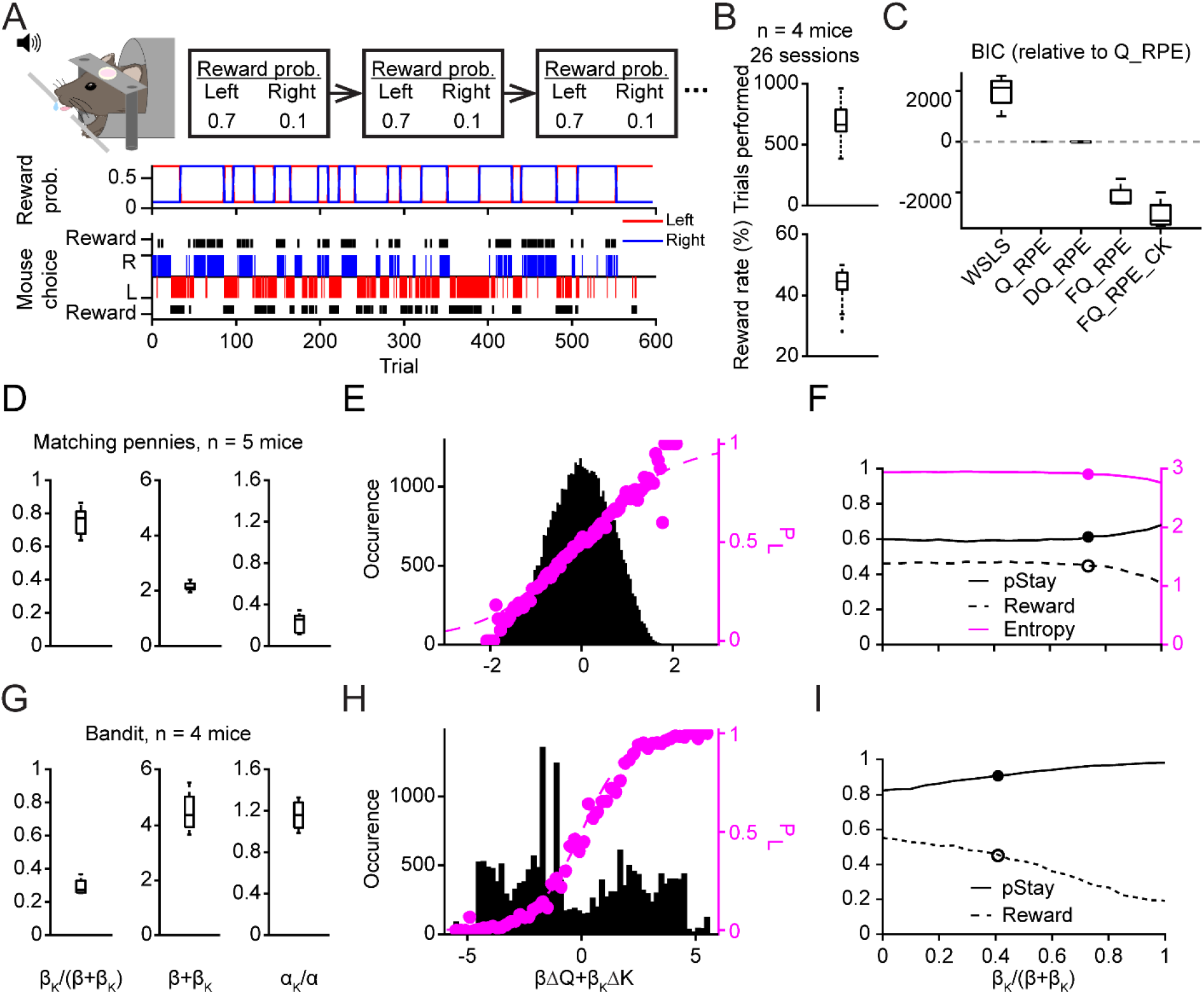
Comparison between matching pennies and two-armed bandit: behavioral results, model fitting, and computer simulations. **(A)** A schematic diagram of the two-armed bandit task. Top: Head-fixed mouse makes a left or right choice by licking the spouts. The trials are separated into different blocks based on the high (0.7) or low (0.1) reward probability assigned to each of the two choices. Middle: The assigned reward probabilities for the left (red) and right (blue) choices in an example session. Bottom: The choices and outcomes in the same session. The reward rate was 52.8%. Blue and red bars indicate right and left choices. Black bars indicate rewards. **(B)** Summary from 26 sessions. Top: the average trials performed each session is 702±26. Bottom: the average reward rate is 43.5%±1.1%. **(C)** Model comparison using BIC. **(D)** Learning parameters extracted from fitting the hybrid model to the matching pennies data. Left: the relative weight of choice kernel, *β*_K_/(*β*+*β*_K_) = 0.75±0.04. Middle: sum of inverse temperature, *β*+*β*_K_ = 2.15±0.07. Right: relative learning rate of the choice kernel. *α*_K_/*α* = 0.22±0.04. **(E)** Psychometric curve based on the whole dataset. Black histograms indicate the distribution of trials according to the weighted sum of the difference of action values (Δ*Q*) and the difference of choice kernels (Δ*K*). Dashed purple line shows the predicted probability to choose left according to the softmax equation used by the hybrid model; Purple dots show the observed probability to choose left. **(F)** The performance of a computational agent playing the game, where *β*_K_/(*β*+*β*_K_) was varied, while the learning rates and (*β*+*β*_K_) were fixed and set to be the median of the fitted values based on animal data. The solid and open dots indicate the median *β*_K_/(*β*+*β*_K_) value fitted based on animal data. **(G - I)** Same as (D - F) for the two-armed bandit task.

A head-to-head comparison of the fitted learning parameters highlights the distinct modes of operation employed by the mice for matching pennies versus two-armed bandit (***Figure 3D – I***). In matching pennies, the choice kernel was weighed more strongly than reinforcement learning during action selection (*β*_K_/(*β*_K_+*β*) = 0.76±0.04; ***Figure 3D***), although the sum of inverse temperatures is relatively low, indicating high level of exploration (*β*_K_+*β* = 2.15± 0.07). The ratio of learning rates suggests that the choice kernels were updated slower than the action values (*α*_K_/*α* = 0.22±0.04), which is illustrated by the example in ***Figure 2C***, where the action values fluctuate more rapidly than the choice kernels. In two-armed bandit, the fitted learning parameters were significantly different. Namely, choice kernels had less weight (*β*_K_/(*β*_K_+*β*) = 0.29±0.03; ***Figure 3G***; P = 0.02, two-sided Wilcoxon rank-sum test) and inverse temperature was higher (*β*_K_+*β* = 4.50±0.40; P = 0.02), indicating that the animals relied more on reinforcement feedback to guide decisions and had a lower tendency to explore.

These disparate sets of fitted model parameters for the two tasks led to learning in different regimes, which could be visualized by looking at the weighted sum of action-value and choice-kernel differences, the crucial parameter for the softmax function for action selection each trial. Here, on most matching pennies trials, animals decided with a weighted sum close to zero, i.e., with near equal probabilities of choosing left and right, making it difficult for the computer opponent to predict their choice (***Figure 3E***). By contrast, performance during two-armed bandit involved weighted sums lying at more extreme values (***Figure 3H***). This difference is consistent with the animal spending considerable number of trials exploiting the high-reward-probability side in a block, and only needed to adapt around a block switch. In both cases, the observed choice probability (dots) fitted well to the softmax function used for action selection (dashed line, ***Figure 3E and 3H***).

Finally, to determine how varying the balance between reinforcement learning and choice kernel would affect performance, we simulated choice behavior in the two tasks using a computer agent, varying *β*_K_/(*β*_K_+*β*) while fixing *α*_K_/*α* and *β*_K_+*β*. For matching pennies, the simulations revealed that the reward rate was relatively stable if the computer agent used mostly reinforcement learning or a hybrid strategy (***Figure 3F***). However, if the computer agent based its actions exclusively on choice kernels, the performance declined precipitously. It was intriguingly to see the *β*_K_/(*β*_K_+*β*) estimated from behavioral data lied around the threshold between these two conditions, indicating that the animals might have settled on a hybrid strategy that balanced a tradeoff between reward and effort. For two-armed bandit, reward rate is maximized if the agent uses only reinforcement learning to guide decisions. However, we find that the animals have residual tendencies captured by choice kernel and therefore lie away from optimal performance (***Figure 3I***). Collectively, the direct comparison across tasks illustrates the utility of matching pennies for exploring a regime in reinforcement learning that is distinct from the more widely used two-armed bandit paradigm.

### The pupil response contained choice- and outcome-related signals during matching pennies

While the mouse played matching pennies, a camera was positioned to capture a video of the right eye (***Figure 4A***). We used the DeepLabCut toolbox (Mathis et al., 2018; Nath et al., 2019) to extract pupil size from the videos (see Material and Methods). ***Figure 4B*** shows an example frame with manually selected and computer-generated labels. To quantify the quality of the automatic labeling, we calculated the deviations between the manually selected and automatically estimated labels for the uppermost, lowermost, leftmost, rightmost, and central points of the mouse’s pupil (***Figure supplement 2A - B***). The mean values of the deviations were close to zero, demonstrating that the estimates had little bias. To give an intuition into the pupil size fluctuation observed, for one session, we plotted the time course of the pupil diameter after z-score normalization (***Figure supplement 2C***). When aligned to select trial types, there were obvious task-related transients in the pupil size (***Figure supplement 2D***). In this study, we were interested in the relation between pupil fluctuations and decision-related variables on a trial-by-trial basis, therefore we calculated the pupil response for each trial – which was defined as the pupil size in z-score minus the pre-cue baseline z-score (***Figure 4C***).

**Figure 4.**
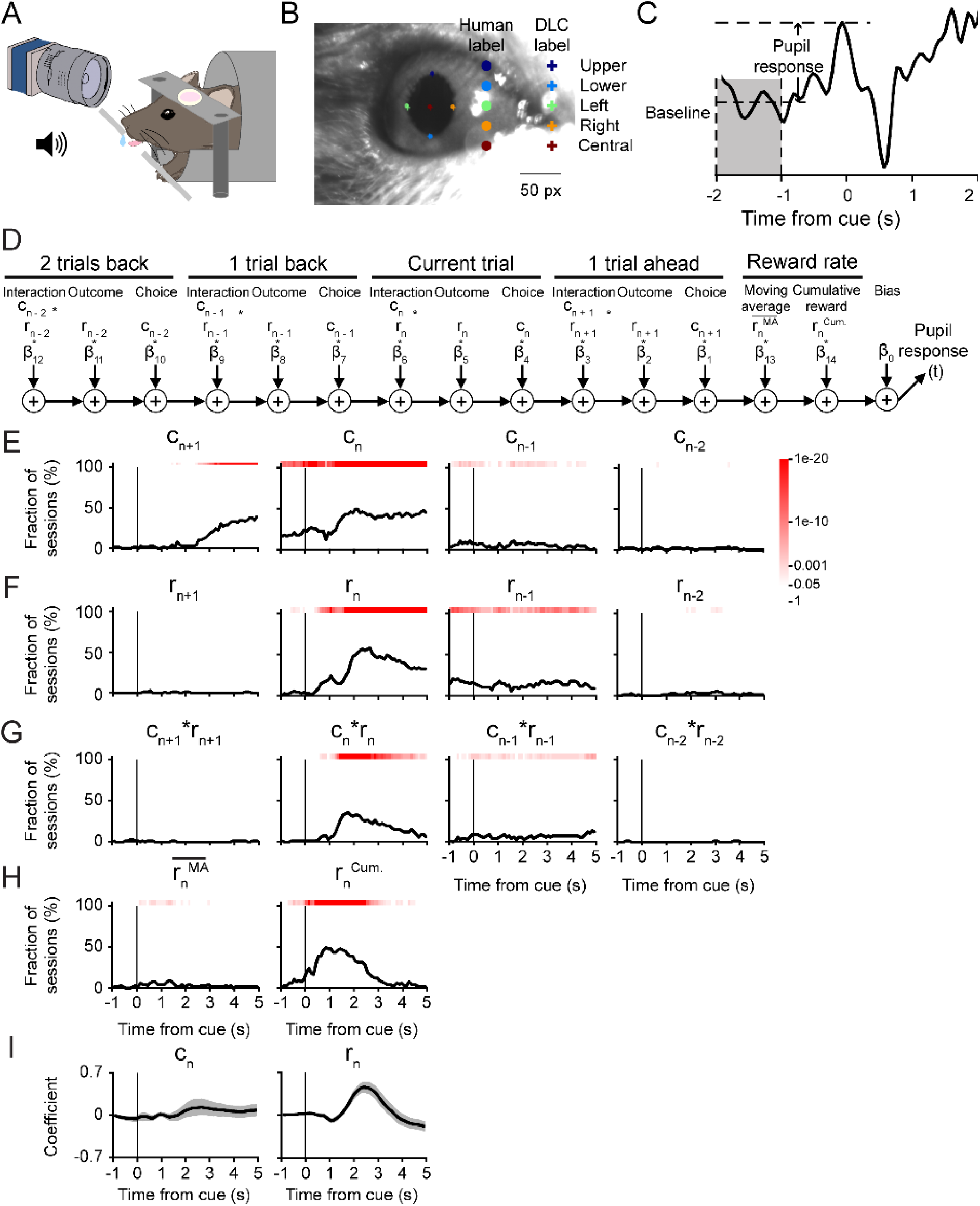
Effects of choices and outcomes on pupil responses during matching pennies. **(A)** A schematic illustration of the pupillometry set up. **(B)** An example still frame from a video showing both human labeling (dot) and DLC labeling (cross) for five labels. Scale bar, 50 pixels. **(C)** The pupil response at any time during a trial (−1 to 5 s from the cue) is the z-score at the corresponding time subtracted by the baseline, which is the mean z-score between −2 and −1 s before the cue onset. **(D)** A schematic diagram of the multiple linear regression model, i.e., equation (9), that was fit to the pupil response in each 100 ms time bin. **(E)** The fraction of sessions with significant regression coefficient for choice in the next trial *c_n+1_*, choice in the current trial *c_n_*, choice in the previous trial *c_n-1_*, and choice in the trial before the previous trial *c_n-2_*. Red shading indicates the p-value from the chi-square test, without correcting for multiple comparison. **(F)** Same as (E) for trial outcomes. **(G)** Same as (E) for the interactions of choice and outcome. **(H)** Same as (E) for recent reward rate, calculated as a moving average over last 20 trials, and the cumulative reward from start of session to current trial. **(I)** The mean regression coefficients of several predictors: choice of the current trial (*c_n_*), reward of the current trial (*r_n_*). Shading indicates the 95% confidence interval estimated by bootstrap.

To characterize the factors that drive pupil responses during matching pennies, we used multiple linear regression. For each session, we fitted a regression model to determine the relation between the pupil responses and the choices, outcomes, reinforcers (choice-outcome interactions), the recent reward rate, and the cumulative reward (***Figure 4D***). Specifically, the choices, outcomes, and interactions included terms for the next trial, the current trial, the last trial, and the trial before last to capture the potential persistent effects of these variables on neural correlates (Bari et al., 2019; Seo and Lee, 2007; Siniscalchi et al., 2019; Sul et al., 2010). The analyses revealed that pupil responses were modulated by choices, outcomes, and reinforcers during matching pennies (***Figure 4E – G***). For a significant fraction of sessions, we detected a change in pupil size signaling the upcoming choice well before the trial would start (*C_n+1_*, ***Figure 4E***). The early choice-related signal suggested that the animal was planning and preparing for the upcoming action prior to the cue. The choice-related signal ceased abruptly at around the time of the cue onset for the next trial (*C_n-1_*, ***Figure 4E***). Meanwhile, outcome- and reinforcer-related signals in the pupil responses emerged after the potential reward would be delivered and persisted for the next two trials (*r_n-1_, r_n-2_, c_n-1_ * r_n-1_, c_n-2_ * r_n-2_*, ***Figure 4F, G***). The pupil responses were also influenced by the cumulative reward (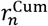, ***Figure 4H***), which related to the number of trials performed, and therefore might reflect the motivational state of the animal. Moreover, although choice consistently influenced the pupil responses, the mean amplitude of the effect was more muted than outcome. Specifically, the presence of a reward led a large phasic dilation of the pupil (*r_n_*, ***Figure 4I***).

### Pupil response was modulated by latent variables related to value updating, but not action selection

Previously we showed that the animals’ behavior can be captured by the hybrid FQ_RPE_CK model, therefore we next asked whether pupil responses may accompany changes in select latent decision variables. To this end, we built additional multiple linear regression models using latent variables relevant for action selection or value updating. For action selection, we considered action-value difference, chosen value, choice-kernel difference, and chosen choice kernel (***Figure 5A***). Consistent with our prior observation, we found significant choice- and outcome-related signals (***Figure 5B - C***), however there was no reliable, sustained modulation of the pupil responses by the latent variables used for action selection (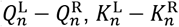 ***Figure 5D - E***). For value updating, we tested latent variables including reward prediction error (RPE) and choice kernel error (CKE) in addition to action-value difference and choice-kernel difference (***Figure 6A***). In a significant fraction of sessions, pupil responses were modulated by the RPE, and to a lesser extent CKE and choice-kernel difference (***Figure 6B – D***). Characteristically, mean coefficient was positive for both trials involving positive or negative RPE (***Figure 6E***). In other words, positive RPE led to phasic dilation of the pupil, whereas negative RPE was associated with transient reduction in pupil size. Overall, these analyses show that transient change in pupil diameter is modulated by latent variables used in value updating, namely the RPE.

**Figure 5.**
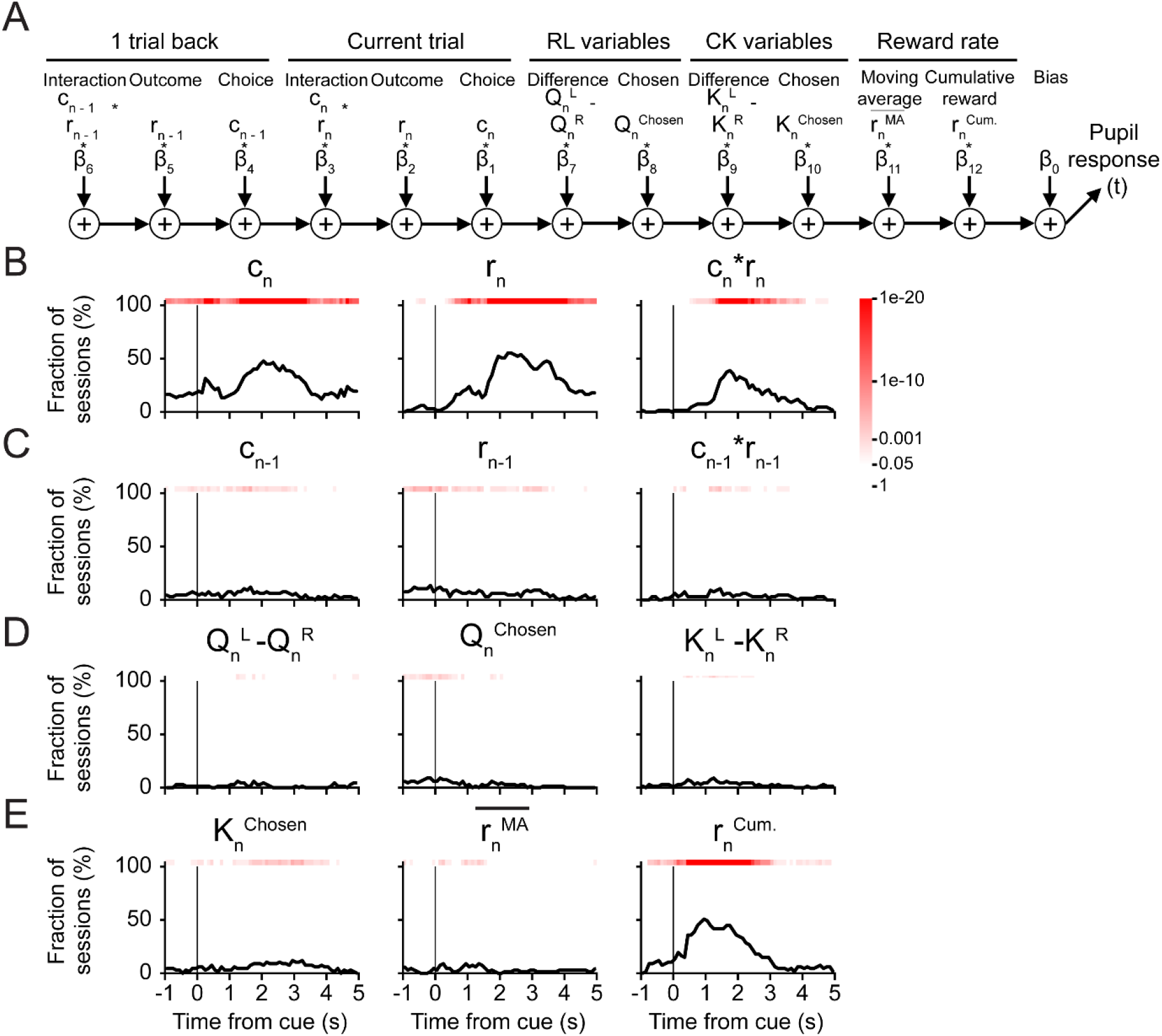
Effects of decision variables for action selection on pupil responses during matching pennies. **(A)** A schematic diagram of the multiple linear regression model, i.e., equation (12). **(B)** The fraction of sessions with significant regression coefficient for the choice *c_n_*, outcome *r_n_*, and the interaction *c_n_* x *r_n_* in the current trial. Red shading indicates the p-value from the chi-square test, without correcting for multiple comparison. **(C)** Same as (B) for the choice, outcome, and interaction in the previous trial. **(D)** Same as (B) for the difference in action values 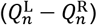, the action value of the chosen action 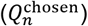, and the difference in the choice kernel 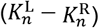. **(E)** Same as (B) for choice kernel of the chosen action 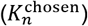, moving-average reward rate 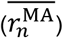, and cumulative reward 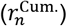.

**Figure 6.**
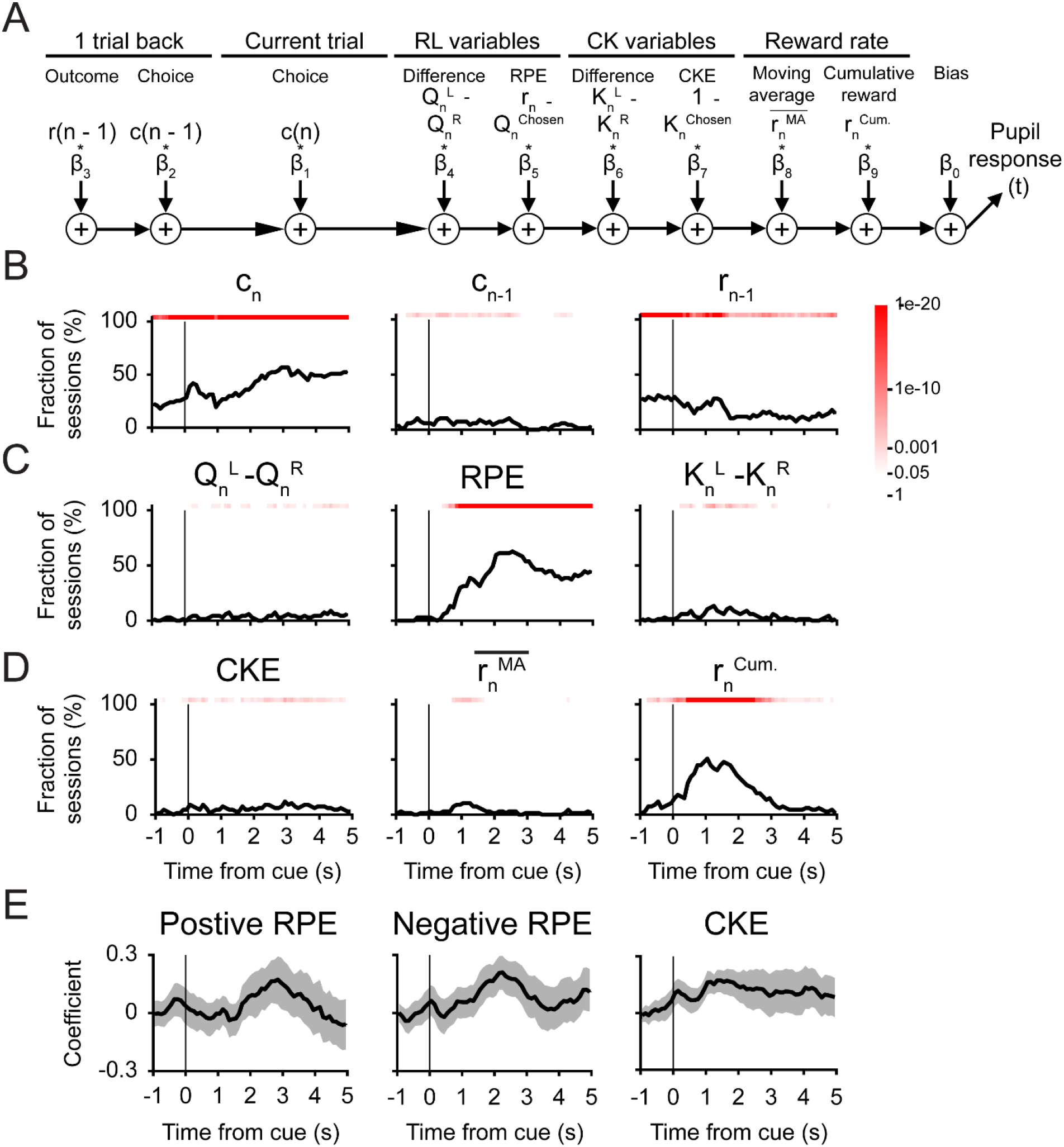
Effects of decision variables for value updating on pupil responses during matching pennies. **(A)** A schematic diagram of the multiple linear regression model, i.e., equation (13). **(B)** The fraction of sessions with significant regression coefficient for choice in the current trial *c_n_*, choice in the previous trial *c_n-1_*, and outcome in the previous trial *r_n-1_*. Red shading indicates the p-value from the chi-square test, without correcting for multiple comparison. **(C)** Same as (B) for difference in action values 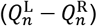, the RPE, and the difference in choice kernels 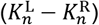. **(D)** Same as (B) for the choice kernel error (*CKE*), moving-average reward rate 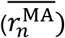, and cumulative reward 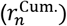. **(E)** The mean regression coefficients for RPE (quantified separately for trials with positive or negative RPE) and CKE. Shading indicates the 95% confidence interval estimated by bootstrap.

### Similar pupil correlates for decision-related variables during two-armed bandit task

The results presented so far indicated that for mice playing the matching pennies game, the transient pupil dilations were modulated by choice, outcome, and latent decision variables relevant for value updating. To determine if comparable pupil responses occur for two-armed bandit, we made recordings during the task and applied the same multiple linear regression analyses. We found that the choice- and outcome-dependent signals were present in a significant fraction of sessions (***Figure 7A***). The latent variables for action selection were largely absent (***Figure 7B***). The value updating variables, especially the RPE and again to a lesser degree the CKE, were associated with transient pupil responses (***Figure 7C***). Therefore, our results show that factors that influenced pupil responses in matching pennies – choice, outcome, and value updating variables – were also contributors to the pupil responses during the two-armed bandit task.

**Figure 7.**
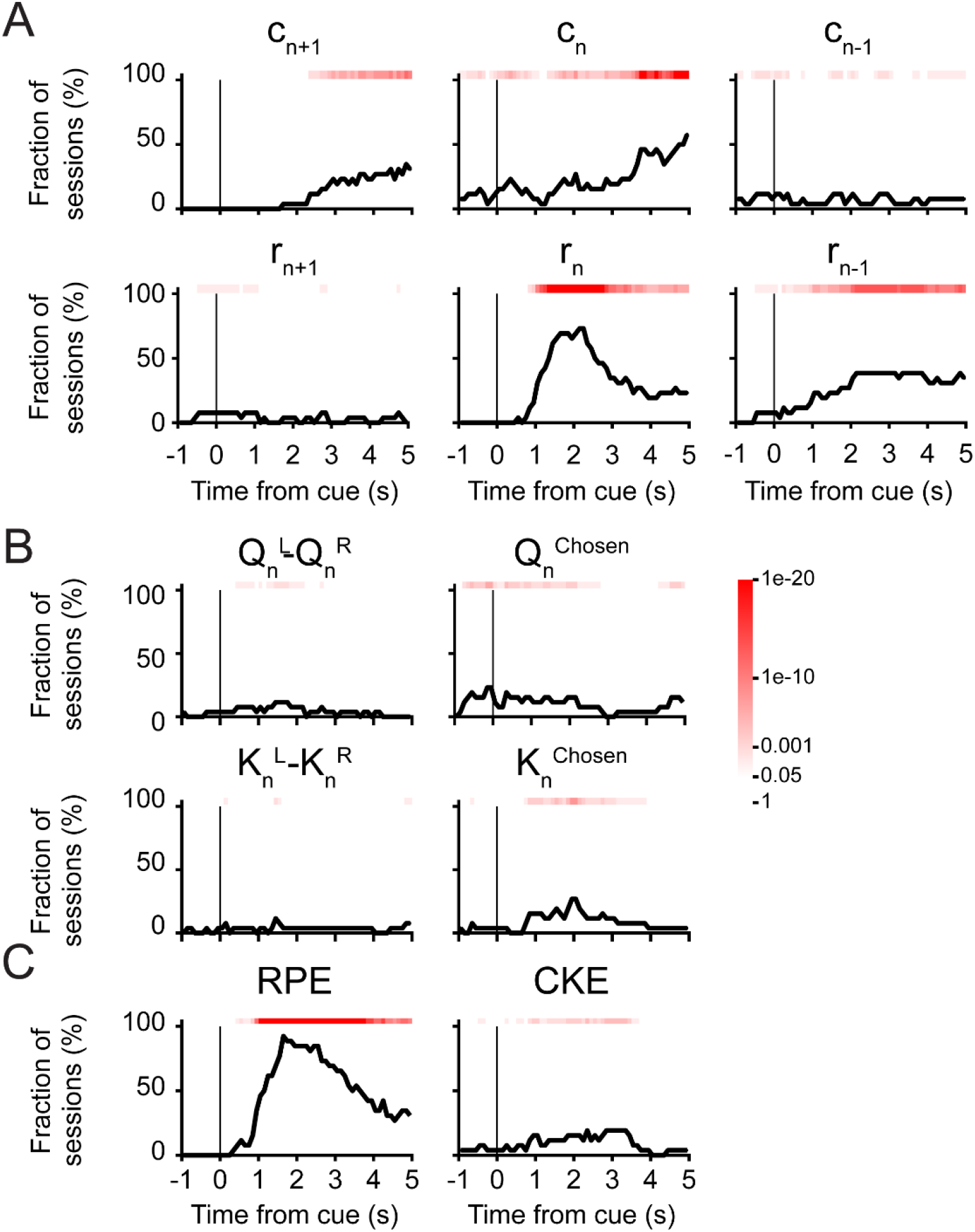
Multiple linear regression analyses of factors influencing pupil responses during two-armed bandit. (A) The fraction of sessions with significant coefficient for choice in the next trial *c_n+1_*, choice in the current trial *c_n_*, choice in the previous trial *c_n-1_*, outcome in the next trial *r_n+1_*, outcome in the current trial *r_n_*, and outcome in the previous trial *r_n-1_*,. The results were obtained by fitting equation (9) as we did in Figure 4 but onto pupil responses from two-armed bandit task. Red shading indicates the p-value from the chi-square test, without correcting for multiple comparison. (B) Same as (A) but for the difference in action values 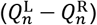, the action value of the chosen action 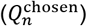, the difference in the choice kernel 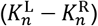, and the choice kernel of the chosen action 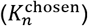. The results were obtained by fitting equation (12) as we did in Figure 5 but onto pupil responses from two-armed bandit task. (C) same as (A) but for the RPE and the choice kernel error (CKE). The results were obtained by fitting equation (13) as we did in Figure 6 but onto pupil responses from two-armed bandit task.

### Correlated fluctuations of left and right pupils during the two-armed bandit task

One intriguing result is that pupil response was influenced by choice, i.e., whether the animal was choosing left or right. Is this a genuine choice-related signal? Some previous studies show that pupil dilations can predict the upcoming choice of human subjects including the choice timing and the selection of one out of five digits (Einhauser et al., 2010), and the decision of a yes/no question (de Gee et al., 2014). However, another possibility is that when the animal made a choice, the tongue lick movement could be associated with facial movements leading to spurious detection of pupil size changes. To clarify the issue, we positioned two cameras to record both eyes simultaneously during the two-armed bandit task (***Figure 8A***). For each eye, we applied multiple linear regression (***Figure 4D***) to analyze the influences of choices and outcomes on pupil responses. We reasoned that if the choice-related signal was a movement artifact, then the aberrant signal should differ across eyes and across animals, and therefore the coefficients extracted from left and right eye would be uncorrelated. By contrast, if the choice-related signal was related to the animal’s internal decision, we would expect consistent dilation responses in both eyes. We analyzed the coefficients for the current choice between 3 – 5 s from cue onset when the pupil responses were largest (***Figure 8B***). The choice-related signal for the left pupil were correlated with that for the right pupil in every session (r = 0.83, *P* = 9 x 10^-43^). The positive correlation coefficient indicated that the pupil size changes are symmetric in the two pupils. As a positive control, we plotted the coefficients of the current outcome in the same time window, which was expected not to be lateralized and indeed showed a positive correlation coefficient (r = 0.90, *P* = 0; ***Figure 8C***). Taken together, these results suggested that the effect of choice on transient pupil response could not be explained by simple movement artifact.

**Figure 8.**
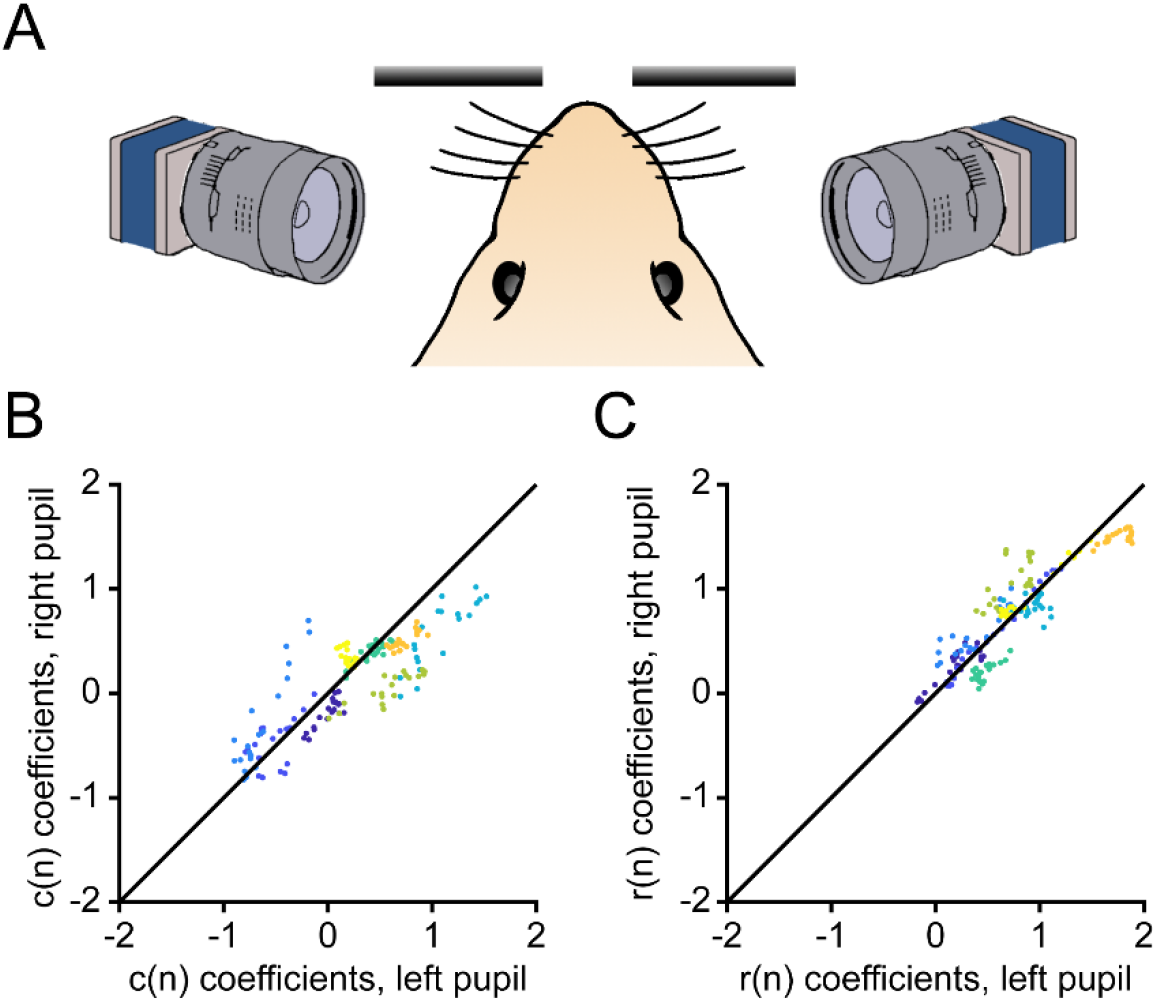
Two-pupil recordings during the bandit task. **(A)** Schematics of the two-pupil recordings setup. Two cameras were placed in front of the two pupils with the same angle while the mouse was performing the task. **(B)** Scatter plot of the linear regression coefficients of the current choice within the 3-5 s from the cue time. The linear regression is the same as shown in Figure 4. X-axis: coefficients of the left pupil; Y-axis: coefficients of the right pupil. Different colors represent different subjects. Each dot represents a 0.1-s interval within the 3-5 s period. The black line shows the diagonal when coefficients of the left pupil equal to that of the right pupil. **(C)** Same as (B) for current outcome.

## DISCUSSION

The present study has two main findings. First, we demonstrated that head-fixed mice can play a two-player competitive game against a computer opponent. Their tendencies in the matching pennies game can be described by a computational model incorporating reinforcement learning and choice kernels. Second, we showed that transient pupil responses of the animals were associated with observable variables such as choices and outcomes, as well as latent variables relevant for value updating, but not action selection.

### Performance in the matching pennies game

Iterative matching pennies is a classic competitive game. Subjects playing the game tend to deviate from the Nash Equilibrium. For example, human players attempting to generate random choices would switch too often (Camerer, 2003). Macaque monkeys and chimpanzees also showed deviation from the optimal strategy (Lee et al., 2004; Martin et al., 2014; Seo et al., 2009; Seo and Lee, 2009; Soltani et al., 2006). Rats were shown to counter-predict the opponent’s choice first and switch to a more random behavior when facing a strong competitor (Tervo et al., 2014). Moreover, pigeons playing the matching pennies game exhibit a similar divergence from the optimal play as the humans do (Sanabria and Thrailkill, 2009). Here we showed that head-fixed mouse can play matching pennies at a high level, albeit unsurprisingly also gaining rewards at below optimal rates. Demonstrating that mice can be studied using competitive game paradigms opens new avenues for studying neural circuitry for reward learning in animal models of neuropsychiatric disorders (Barthas et al., 2020; Liao and Kwan, 2021).

Although initial characterizations of the tendencies for sub-optimal play have relied on standard learning algorithms (Erev and Roth, 1998; Lee et al., 2004), recent studies have reported deviations from the predictions of reinforcement learning (Hampton et al., 2008; Seo et al., 2014; Tervo et al., 2014). In this study, we employed a hybrid model, which combines Q-learning with choice kernels, also known as the choice-autocorrelation factor (Katahira, 2015; Wilson and Collins, 2019). Specifically, choice kernels were included to capture serial choice dependency, which is commonly observed in humans and animals performing various tasks (Abrahamyan et al., 2016; Akaishi et al., 2014). The hybrid model indeed fit the behavior better than variations of Q-learning algorithms (***Figure 2B***). We want to highlight the numerical simulations where the reward rate was examined as a function of relative inverse temperature (***Figure 3F***). The animal’s performance lied at a regime where further increase in reliance on choice kernel would deteriorate sharply the reward rate. The result hints at the possibility that the animal may be maximizing a reward-effort trade-off, because repeating the same choice is likely to be less effortful and indeed is the strategy taken by the animal often near the end of a session.

Many studies of flexible decision-making in rodents have relied on two-armed bandit tasks with a block-based design (Bari et al., 2019; Groman et al., 2019; Hattori et al., 2019; Ito and Doya, 2009; Sul et al., 2011; Sul et al., 2010; Tai et al., 2012). This paradigm has many merits, but also a few shortcomings. First, the length of the blocks is a key parameter that defines the volatility of the environment, but is typically manually set by the experimenter. By contrast, matching pennies refers to a payoff matrix for gameplay, but contain no other experimenter-defined parameters. Second, within a block, it is advantageous for the animals to continually exploit the high-reward-probability option, therefore overtraining in the two-armed bandit task may lead to strong serial choice dependencies. To the contrary, in matching pennies, the computer opponent is designed to detect such dependencies and exert competitive pressure on the animal, therefore the animal is encouraged to always diversify its choice patterns during the session. Consequently, two-player games such as matching pennies are elegant and simple in design, and allows for investigation of neural mechanism underlying flexible decision-making under a regime that are quite different from multi-armed bandit tasks (***Figure 3D - I***).

### Pupil responses and potential neuromodulatory mechanisms

Pupil fluctuation is an indicator of the arousal state of an animal, and likely associates with the levels of various neuromodulators in the forebrain (McGinley et al., 2015). The relationship between pupil size and NE is supported by prior results, which showed reliable tracking of pupil fluctuations to activity of noradrenergic axons in the neocortex (Reimer et al., 2016). Furthermore, studies of the locus coeruleus (LC), the main source of NE for the forebrain, demonstrated a correlation between single unit firing in LC and pupil diameter (Aston-Jones and Cohen, 2005; Yang et al., 2021), and pupil dilation triggered by electrical microstimulation of the locus coeruleus (Joshi et al., 2016). Several studies have linked pupil dynamics and activity in LC to choice behavior, such as in the consolidation of the previous choices (Clayton et al., 2004; Einhauser et al., 2010) or the shaping of upcoming actions (de Gee et al., 2014). However, these studies were based on visual perceptual tasks, which is different from our task design where the cue is auditory and carries no relevant information except for trial timing. Our results hint at a potential role for noradrenergic signaling in post-decisional value updating, because the pupil response was correlated with the RPE and weakly associated with choice kernel error. This would agree with work that have showed the importance for rewards in LC activity and pupil dilation in various behavioral settings (Aston-Jones et al., 1997; Sara and Segal, 1991; Varazzani et al., 2015). Furthermore, there is strong evidence linking pupil changes to errors and adaptations in decision tasks involving trial- or block-based inference (Nassar et al., 2012; Urai et al., 2017), highlighting the role of pupil-linked systems to control the influence of incoming data to guide future decisions.

Overall, the current study lays the groundwork for studying reward-based learning in mice using competitive games. The two-player matching pennies game, which we showed the mouse can play against a computer opponent, may potentially be extended to a mouse competing against another mouse in the future. The paradigm may therefore provide a quantitative framework for evaluating social decision-making in mice. The findings of significant pupil correlates to the major decision-related variables during matching pennies provide clues to the neuromodulatory mechanisms in mice. The current results open avenues for future research into the role of neuromodulators in mediating the adaptive and social aspects of decision-making in mice.

## Acknowledgements

We thank Daeyeol Lee and Hyojung Seo for helping with programming the matching pennies game, Michael Siniscalchi on behavioral training, and Matthew McGinley on the pupil recording setup. This work was supported by NIH/NIMH grants R01MH112750 (A.C.K.), R01MH121848 (A.C.K.), R21MH118596 (A.C.K.), China Scholarship Council-Yale World Scholars Fellowship (H.W.), Gruber Science Fellowship (H.K.O.), NIH training grant T32NS007224 (H.K.O.), and Kavli Institute for Neuroscience Postdoctoral Fellowship (H.A.).

## Author Contributions

H.W. and A.C.K. designed the research. H.W. performed the matching pennies experiments. H.K.O. and C.E.M. performed the two-armed bandit experiments. H.W. analyzed the data. H.A. assisted by developing the bandit task and providing code for analysis. H.W. and A.C.K. wrote the paper with input from all other authors.

## Supplementary figures

**Figure supplement 1.**
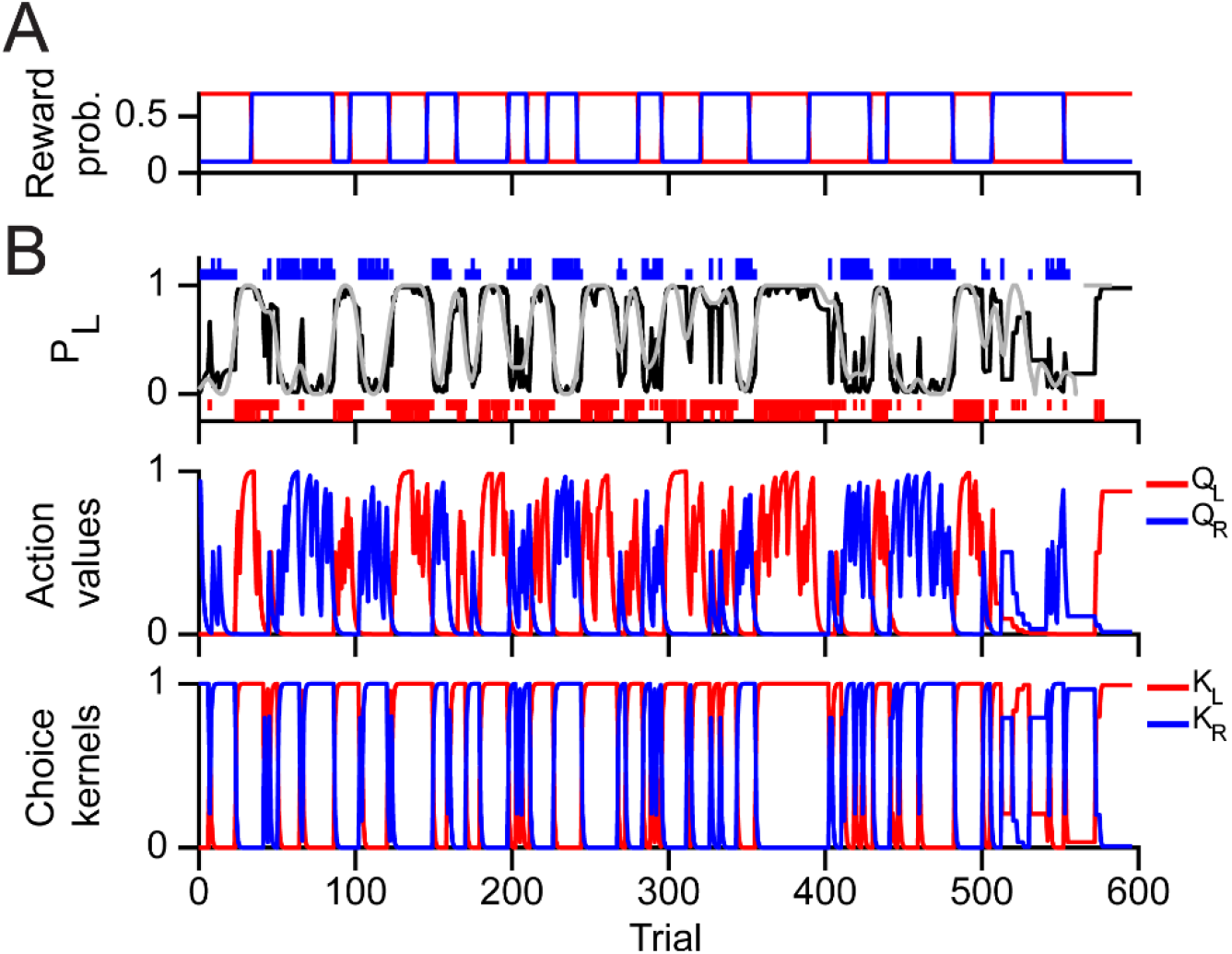
Computational modeling of the animals’ behavior during two-armed bandit. **(A)** The switch of the reward probabilities of left (red) and right (blue) choices. **(B)** An example of the time course of the latent variables and predicted behavior in the same session as (A). Top: Long red and blue bars indicate rewarded left and right choices. Short red and blue bars indicate unrewarded left and right choices. Gray line shows the observed probability to choose left, smoothed by a gaussian kernel. Black line shows the probability to choose left predicted by the hybrid model. Middle: the action values of left (red) and right (blue) choices estimated by the hybrid model. Bottom: the choice kernels of left (red) and right (blue) choices estimated by the hybrid model.

**Figure supplement 2.**
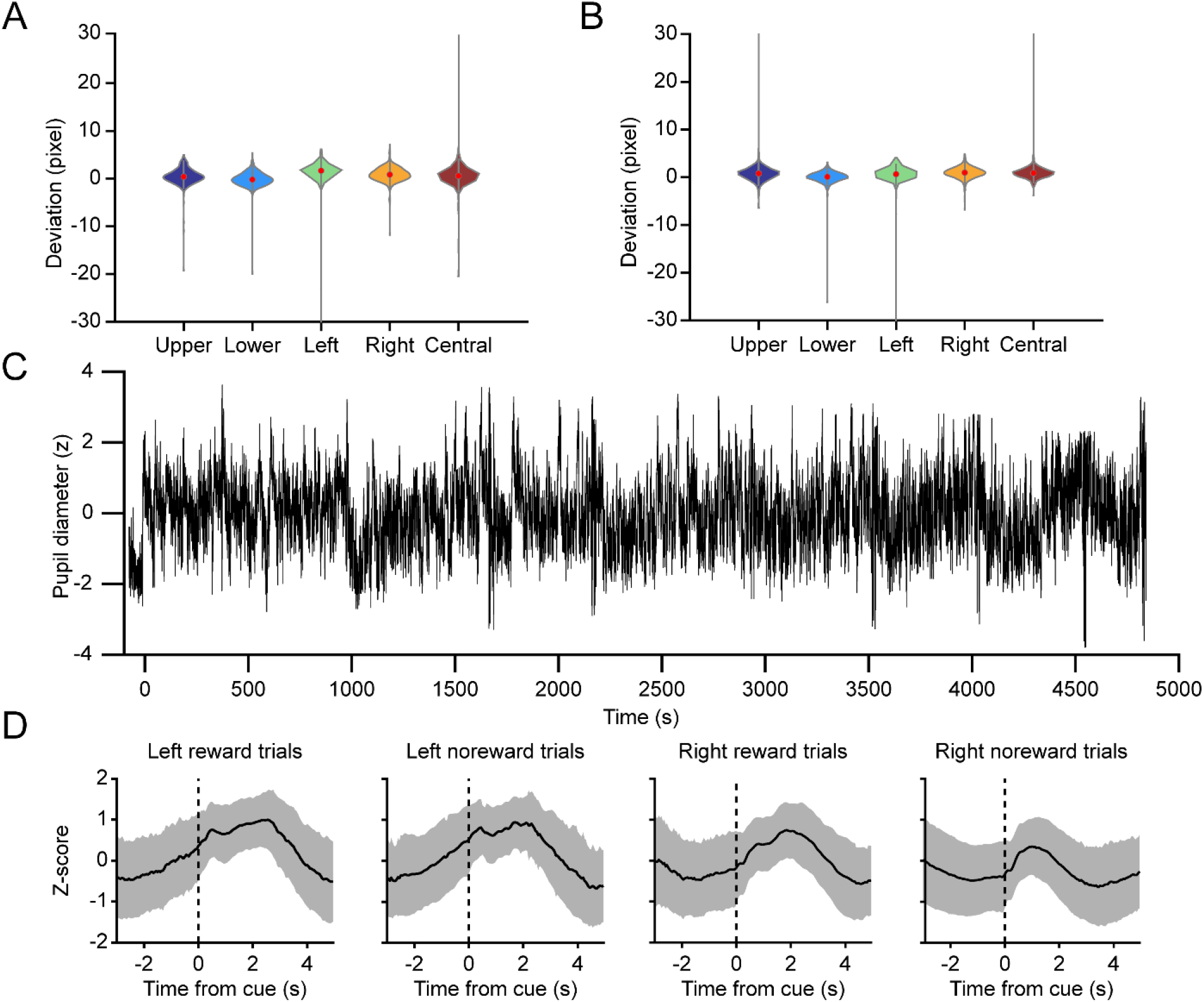
Extracting the pupil size using DLC. **(A)** Deviation of DLC labels from manually selected labels in the x-axis (n = 324 frames, taken from different sessions of multiple animals). **(B)** Same as (A) for y-axis. **(C)** An example trace of the z-score of pupil diameter during a session of matching pennies. **(D)** Mean traces of z-score of pupil diameter for four different trial types: rewarded left choice, unrewarded left choice, rewarded right choice, unrewarded right choice. Shading indicates the standard deviation.

